# Cleavage of the Jaw1 C-terminal region enhances its augmentative effect on the Ca^2+^ release via inositol 1,4,5-trisphosphate receptors

**DOI:** 10.1101/2022.12.10.519934

**Authors:** Takuma Kozono, Chifuyu Jogano, Wataru Okumura, Hiroyuki Sato, Hitomi Matsui, Tsubasa Takagi, Nobuaki Okumura, Toshifumi Takao, Takashi Tonozuka, Atsushi Nishikawa

## Abstract

Jaw1, a tail-anchored protein with 39 carboxyl (C)-terminal amino acids, is oriented to the lumen of the endoplasmic reticulum and outer nuclear membrane. We previously reported that Jaw1, as a member of the KASH protein family, plays a role in maintaining nuclear shape via its C-terminal region. Furthermore, we recently reported that Jaw1 functions as an augmentative effector of Ca^2+^ release from the endoplasmic reticulum by interacting with the inositol 1,4,5-trisphosphate receptors (IP_3_Rs). Intriguingly, the C-terminal region is partially cleaved, meaning that Jaw1 exists in the cell in at least two forms: uncleaved and cleaved. However, the mechanism of the cleavage event and its physiological significance remain to be determined. In this study, we demonstrate that the C-terminal region of Jaw1 is cleaved after its insertion by the signal peptidase complex (SPC). Particularly, our results indicate that the SPC with the catalytic subunit SEC11A, but not SEC11C, specifically cleaves Jaw1. Furthermore, using a mutant with a deficit in the cleavage event, we demonstrate that the cleavage event enhances the augmentative effect of Jaw1 on the Ca^2+^ release ability of IP_3_Rs.

**Summary statement:** The C-terminal region of Jaw1, a tail-anchored protein, is cleaved by signal peptidase complex and this cleavage event enhances the augmentative effect of Jaw1 on the Ca^2+^ release activity of inositol 1,4,5-trisphosphate receptors

## Introduction

More than 400 tail-anchored (TA) proteins, membrane proteins with a single transmembrane domain close to the carboxyl (C)-terminal region, are present in human cells (Kalbfleisch *et al*., 2007). Previous studies have revealed that the biogenesis and insertion into the membrane of TA proteins are mediated by multiple pathways (Aviram *et al*., 2016; Guna *et al*., 2018; Haßdenteufel *et al*., 2017; Schuldiner *et al*., 2008; Stefanovic and Hegde, 2007). The TA proteins targeted to each membrane compartment, including the endoplasmic reticulum (ER), nuclear envelope (NE), mitochondria, Golgi apparatus, and plasma membrane, have various cellular functions, such as membrane trafficking, protein degradation, signal transduction, and the morphological maintenance of organelles.

The linker of nucleoskeleton and cytoskeleton (LINC) complex, which bridges the nuclear double membrane, is a key physical regulator of nuclear shape, position, and dynamics (Cain *et al*., 2018; Lüke *et al*., 2008; Starr, 2009). The LINC complex includes Klarsicht/ANC-1/Syne/homology (KASH) proteins, TA proteins localized at the outer nuclear membrane, and Sad-1/UNC-84 (SUN) proteins located at the inner nuclear membrane. Currently, six KASH proteins and four SUN proteins have been determined in mammals (Kozono *et al*., 2018; Morimoto *et al*., 2012; Starr, 2009). The KASH domain, a carboxyl (C)-terminal region of the KASH protein comprising 20–40 amino acids, interacts with SUN proteins in the perinuclear space, resulting in the formation of the LINC complex (Cruz *et al*., 2020). KASH and SUN proteins are connected with cytoskeletons in the cytosol and with the nuclear lamina in the nucleus; this physical linkage makes it possible to transduce forces between the cytosol and nucleus for nuclear maintenance (Luxton and Starr, 2014; Starr and Fridolfsson, 2010).

Jaw1 is one of the TA proteins localized at the ER and outer nuclear membrane (Behrens *et al*., 1994, Horn *et al*., 2013; Kozono *et al*., 2018). The C-terminal region comprising 39 amino acids is oriented with the ER lumen and perinuclear space. We previously reported that Jaw1 functions as a KASH protein to maintain nuclear shape by interacting with SUN proteins (Kozono *et al*., 2018). Although the KASH domain of Jaw1 is crucial for the interaction with SUN proteins in the perinuclear space (Cruz *et al*., 2020), interestingly, the C-terminal region is partially cleaved after translation, implying the loss of the KASH domain (Behrens *et al*., 1996). Thus, Jaw1 exists in the uncleaved or cleaved forms in cells. However, the cleavage site, the enzymes responsible for cleavage, and the physiological significance of the cleavage remain unknown. Jaw1 is involved in Ca^2+^ release from the ER into the cytosol upon the stimulation of G protein-coupled receptor (GPCR) through the interaction with inositol 1,4,5-trisphosphate receptors (IP_3_Rs), calcium channels on the ER membrane (Chang *et al*., 2021; Okumura *et al*., 2022; Prüschenk *et al*., 2021; Shindo *et al*., 2010). Thus, Jaw1 is a bifunctional protein as a KASH protein and as an IP_3_R regulator; however, the underlying mechanisms of these functions remain unclear.

The signal peptidase complex (SPC) is a eukaryotic signal peptidase found in the ER, and it co- and post-translationally cleaves signal peptides in the amino-terminal region of secretory or membrane proteins (Paetzel *et al*., 2002). In addition, the SPC is responsible for the cleavage of TA proteins, but the underlying mechanisms remain unknown (Nilsson *et al*., 2002). The human SPC exists as a heterotetramer consisting of three accessory subunits and a catalytic subunit (Liaci *et al*., 2021). The three accessory subunits are Signal peptidase complex subunit 1 (SPCS1), SPCS2, and SPCS3, and the catalytic subunit is either SEC11A or SEC11C, thus, SPC exists in two different forms: SPCS1–3 plus SEC11A or SPCS1–3 plus SEC11C. However, the differences in its activities and those of the substrates remain elusive.

In this study, we first aimed to identify the cleavage site of the Jaw1 C-terminal region and identified SPC as the enzyme responsible for the cleavage. Particularly, SPCs with SEC11A but not SEC11C as their catalytic subunits specifically processed the cleavage. Furthermore, on the basis of the identified cleavage site, we successfully created a mutant with a deficit in the cleavage event. Using this mutant, we determined that the cleavage event at the C-terminal region enhances the augmentative effect of Jaw1 on the Ca^2+^ release from the ER via interaction with IP_3_Rs.

## Results

### C-terminal cleavage of Jaw1 is unique among KASH proteins

As mentioned above, six KASH proteins, including Jaw1, have been reported to date (Kozono *et al*., 2018; Morimoto *et al*., 2012; Starr, 2009). As observed in the alignment of the amino acid sequences corresponding to the luminal region of each KASH protein, the luminal region of Jaw1 is relatively unique among KASH proteins in both mice and humans, especially in the amino acids sequence near the transmembrane domain (Fig. 1A,B). We therefore investigated whether the C-terminal cleavage event is unique in Jaw1 among all KASH proteins. To confirm this, chimeric mutants in which the luminal region of mouse Jaw1 (Ms Jaw1) was replaced with that of other KASH proteins were expressed in Flp-In T-REx HEK293 FLAG Ms Jaw1 KASH1–KASH5 (K1–K5) (Fig. 1C), and the band pattern was compared with that of N-terminal HA FLAG tandem-tagged Ms Jaw1 (FLAG Ms Jaw1) by western blotting. Consistent with the previous report (Behrens *et al*., 1996), two bands (hereafter called uncleaved form (upper band) and cleaved form (lower band)) were detected in the lane of FLAG Ms Jaw1 (Fig. 1D,E). Importantly, the single band was detected in all the lanes of KASH chimeras. This result indicates that the C-terminal cleavage event of Jaw1 is unique among KASH proteins. The luminal regions of KASH proteins interact with SUN proteins, thereby the KASH proteins are strongly localized at the NE (Horn *et al*., 2013; Starr, 2009). Meanwhile, Jaw1 is localized at both the NE and the ER, as we previously reported (Kozono *et al*., 2018). To compare the localization between Jaw1 and the chimeric mutants, the cells were immunostained with an anti-FLAG antibody. The confocal images showed that the chimeric mutants were strongly localized at the NE, rather than the ER, compared with FLAG Ms Jaw1 (Fig. 1F), indicating that the C-terminal region of Jaw1 is crucial to maintain the localization at both the NE and the ER.

**Figure 1.**
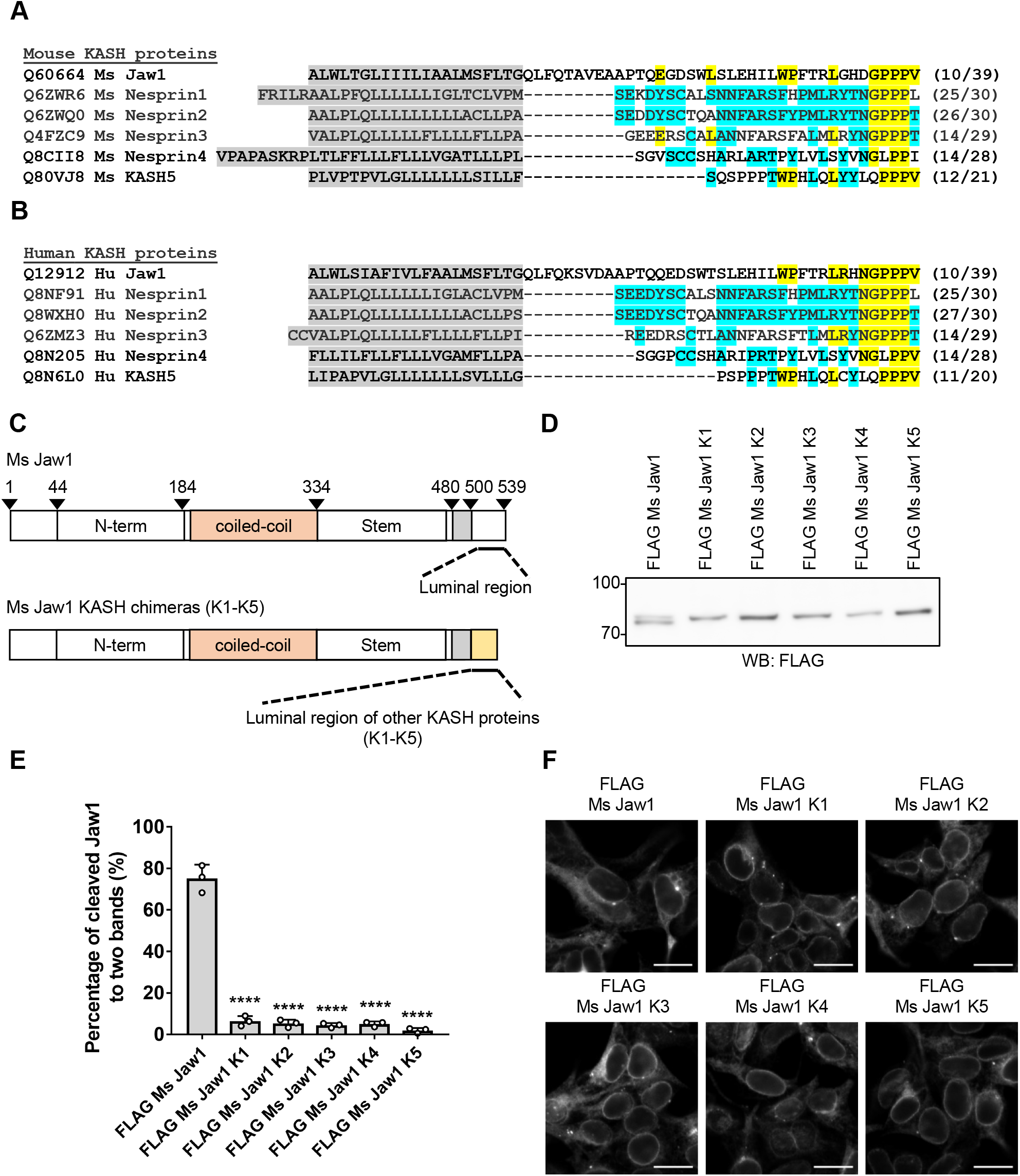
Investigation directed to the uniqueness of the Jaw1 C-terminal cleavage event among KASH proteins. **A, B**) Alignment of amino acid sequences corresponding to the transmembrane domain (gray) and luminal region of Ms Jaw1 (A) and Hu Jaw1 (B) with other KASH proteins. The amino acids conserved between Jaw1 and other KASH proteins are in yellow and those conserved among KASH proteins except Jaw1 are in light blue. The UniProt accession numbers for each gene are shown with the protein name. The number of conserved amino acids in the luminal region with other proteins is shown on the right. **C**) Schematic representation of Ms Jaw1 and Ms Jaw1 KASH chimeras (K1–K5). **D, F)** Flp-In T-REx HEK293 FLAG Ms Jaw1 or FLAG Ms Jaw1 K1-K5 cells were treated with Dox for 24 h. The cells were then subjected to western blotting (D) and immunostaining (F) using an anti-FLAG mouse antibody. Scale bar: 20 μm. **E)** Graph showing the percentage of Jaw1 C-terminal cleavage in (D). The averages of three independent experiments per condition are shown in the graph. Error bar shows ±SD, \*\*\*\**P* < 0.0001; statistical analysis, one-way ANOVA followed by Dunnett’s multiple comparison test.

### C-terminal 30 amino acids are post-translationally cleaved

As mentioned above, although the C-terminal region of Jaw1 has been reported to be cleaved post-translationally (Behrens *et al*., 1996), the mechanism underlying this cleavage event remains unknown. Therefore, we first attempted to identify the cleavage site. To accomplish this, we prepared the plasmids coding for N-terminal HA FLAG tandem-tagged mutants (FLAG Ms Jaw1 504-508A, 507-511A, 512-516A, 517-521A, 522-526A, 527-531A, 532-536A, and 535-539A), in which five sequential residues in the luminal region of Jaw1 were substituted with alanine residues (Fig. 2A). Each mutant was expressed in HEK293 cells and the lysate was subjected to western blotting to compare the band patterns with FLAG Ms Jaw1. However, both the uncleaved and cleaved forms of Jaw1 were detected in all the lanes of mutants, comparable with FLAG Ms Jaw1, although there were subtle differences in the proposition of the two forms among them (Fig. 2B,C). This result indicates that all the mutated sites in the above mutants are not a candidate for the cleavage site. Therefore, we focused on two alanine residues (509/510) that were not substituted in the above mutants. To investigate whether these two alanine residues are crucial for the cleavage event, we prepared the plasmid encoding FLAG Ms AASS, in which these two alanine residues are substituted with serine residues, and compared the band patterns (Fig. 3A). As shown in Figure 3B and 3C, in the lane of FLAG Ms AASS, a single band corresponding to uncleaved Jaw1 was detected, whereas both forms were detected in the lane of FLAG Ms Jaw1. Intriguingly, the luminal regions were highly conserved between mice and humans, especially the two alanine residues (Fig. 3A). Therefore, plasmids encoding N-terminal HA FLAG tandem-tagged human Jaw1 (FLAG Hu Jaw1) or the mutant (FLAG Hu AASS) were prepared and the band patterns were compared. As a result, a single band corresponding to uncleaved Jaw1 was detected (Fig. 3D,E). These results indicate that these alanine residues are essential for the cleavage event of Jaw1. Furthermore, to ensure the cleavage site, we analyzed the amino acid sequence of the cleaved C-terminal fragment using a protein sequencer. Here, we prepared the plasmid encoding Ms Jaw1 NIDR PA, in which the N-terminal and C-terminal regions were fused with the HA FLAG tandem tag and the N-terminal intrinsically disordered region (NIDR) (19.8 kDa) of mouse Jaw1 plus PA tag (1.2 kDa), respectively, which was cleaved into two fragments in the cells (Fig. 3F). The NIDR, a structural flexible region previously described in our study (Kozono *et al*., 2021), was fused between the C-terminal region of Jaw1 and the PA tag to increase the molecular mass of Fragment 2 on the electrophoresis gel. As shown in Figure 3G, Fragment 2 immunoprecipitated by anti-PA beads was specifically detected by SDS-PAGE followed by silver staining (closed black triangle), which was not detected in the lane of Ms Jaw1 PA as a control. Although the band corresponding to Fragment 2 on SDS-PAGE appeared at a higher position than predicted, it is probably due to the low pI derived from NIDR (pI 4.18) and PA tag (pI 3.49). Furthermore, the immunoprecipitated sample was subjected to SDS-PAGE followed by blotting and Coomassie brilliant blue (CBB) staining (Fig. 3H). The amino acid sequence of the band corresponding to Fragment 2 was then analyzed using a protein sequencer. The results highlighted sequence 1 (APTQEGD) in addition to sequence 2 (TQEGDS_LSL) (Figs. 3I,S1,S2). In summary, these data suggest that the C-terminal region of Jaw1 is cleaved between two alanine residues (509/510), although the cleaved peptide might be trimmed more.

**Figure 2.**
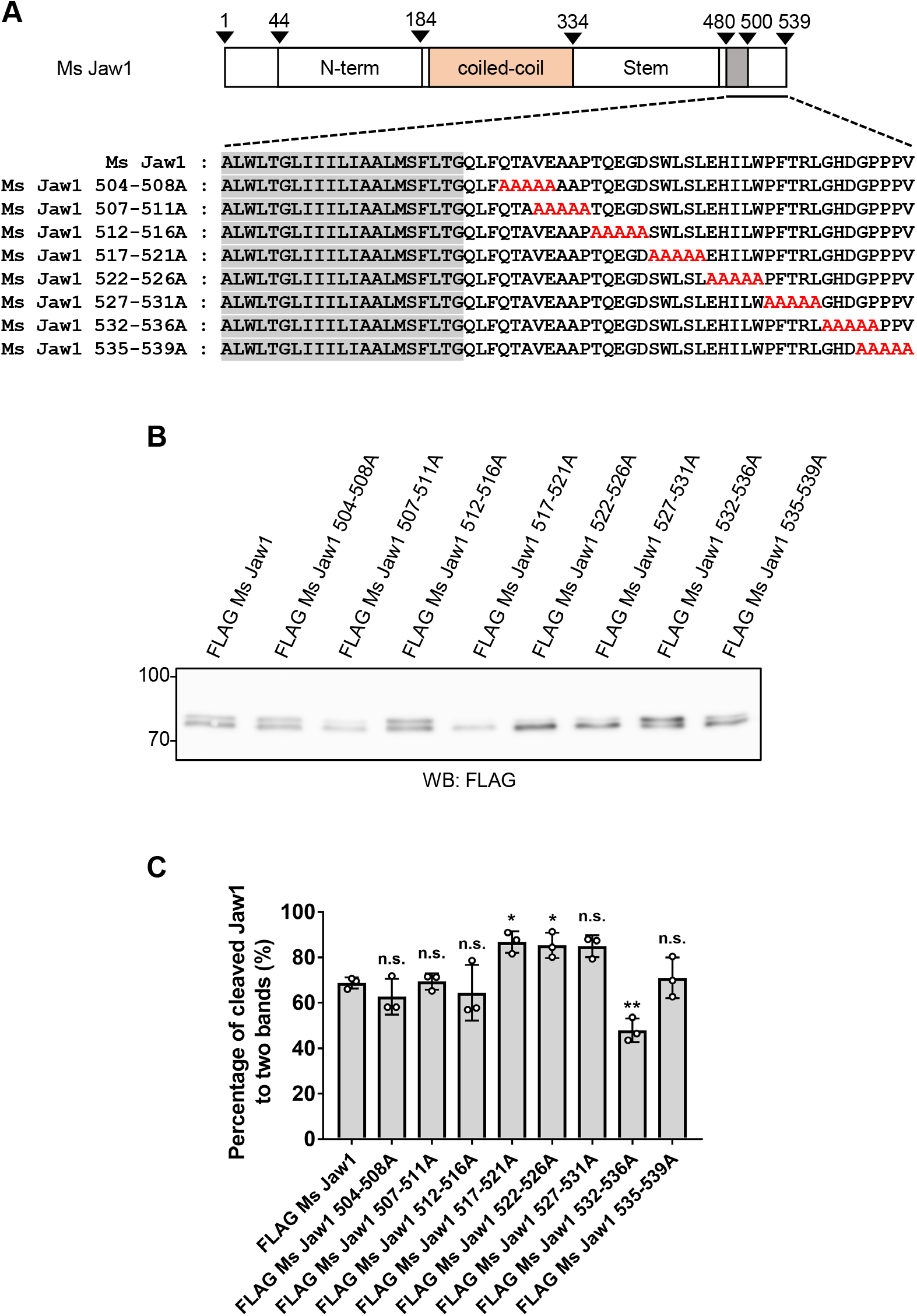
Exploration of Jaw1 C-terminal cleavage site. **A**) Amino acid sequences of mutants in which several sequential amino acids in the luminal region were substituted with alanine residues (red letters). **B**) FLAG Ms Jaw1 and mutants were expressed in HEK293 cells by transfection. After incubation for 24 h, the lysates were subjected to western blotting using an anti-FLAG mouse antibody. **C)** Graph showing the percentage of Jaw1 C-terminal cleavage in (B). The averages of three independent experiments per condition are shown in the graph. Error bar shows ±SD, “n.s.”, not significant; \**P* < 0.05; \*\**P* < 0.01; statistical analysis, one-way ANOVA followed by Dunnett’s multiple comparison test.

**Figure 3.**
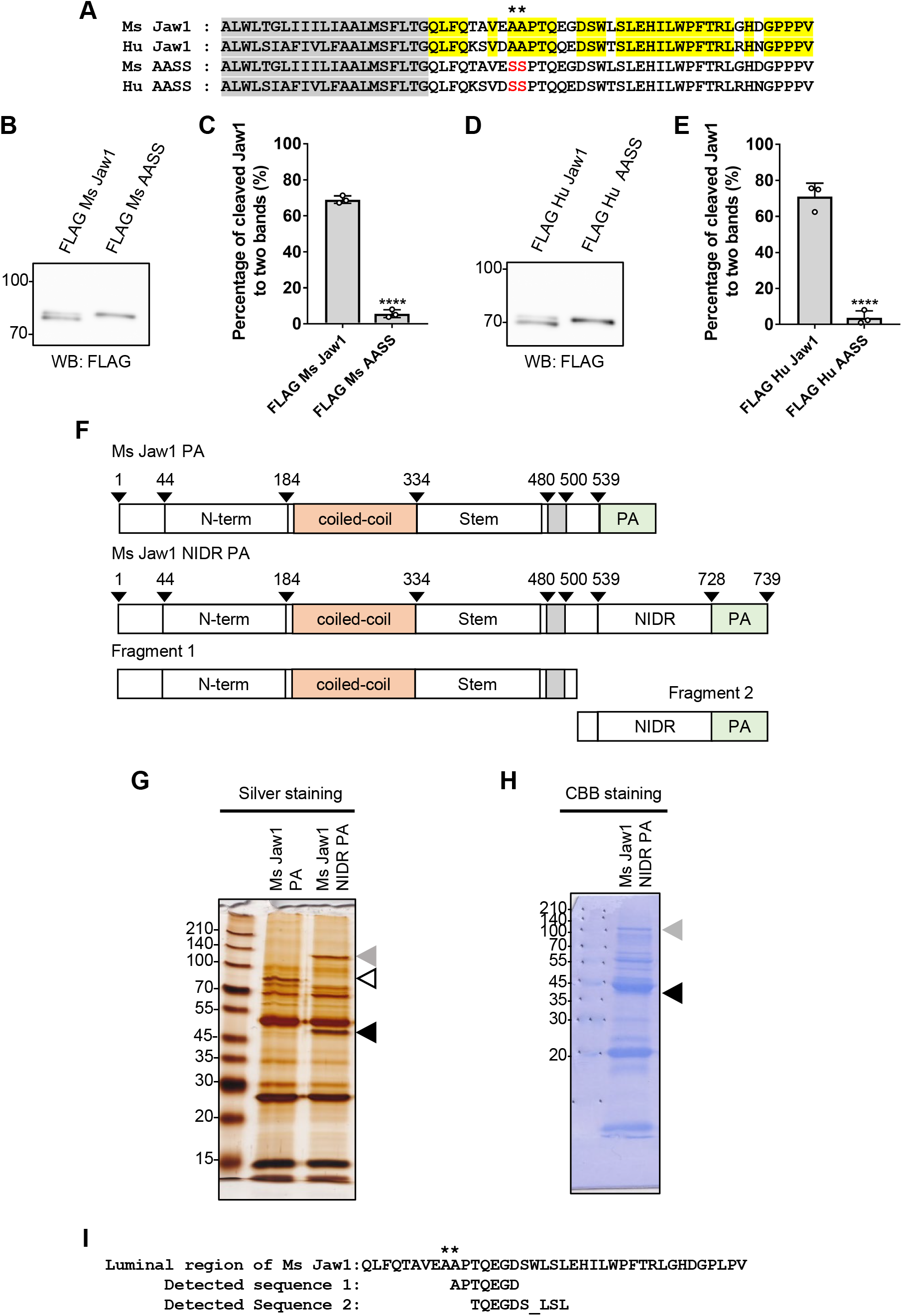
Identification of Jaw1 C-terminal cleavage site. **A**) Alignment of amino acid sequences corresponding to the transmembrane domain (gray) and luminal regions of Ms Jaw1 and Hu Jaw1 and amino acid sequences of AASS mutants. The amino acids conserved in Ms Jaw1 and Hu Jaw1 are in yellow. The mutated sites are in red. Asterisks show the two alanine residues. **B, D**) FLAG Ms Jaw1 and FLAG Ms AASS (B) or FLAG Hu Jaw1 and FLAG Hu AASS (D) were expressed in HEK293 cells by transfection. After incubation for 24 h, the lysates were subjected to western blotting using an anti-FLAG mouse antibody. **C, E)** Graphs showing the percentage of Jaw1 C-terminal cleavage in (B) and (D), respectively. The averages of three independent experiments per condition are shown in the graphs. Error bar shows ±SD, \*\*\*\**P* < 0.0001; statistical analysis, two-tailed Student’s *t*-test. **F**) Schematic representation of Ms Jaw1 PA and Ms Jaw1 NIDR PA. Ms Jaw1 NIDR PA was post-translationally cleaved into Fragments 1 and 2. **G, H**) Ms Jaw1 PA and Ms Jaw1 NIDR PA were expressed in HEK293 cells by transfection. After incubation for 24 h, the lysates were subjected to immunoprecipitation using anti-PA beads. The purified samples were subjected to SDS-PAGE electrophoresis followed by silver staining (G) or blotting and Coomassie brilliant blue (CBB) staining (H). Open triangles, the band of Ms Jaw1 PA; closed triangles, the bands of uncleaved Ms Jaw1 NIDR PA (gray) and Fragment 2 (black). The band corresponding to Fragment 2 in (H) was cut out and subjected to protein sequencing. **I**) The N-terminal amino acid sequences of Fragment 2 detected using protein sequencing. Asterisks show the two alanine residues.

### The SPC cleaves the C-terminal region of Jaw1

To identify the enzyme responsible for the cleavage of the Jaw1 C-terminal region, we focused on the SPC, which was discussed as a candidate in a previous report (Behrens *et al*., 1996). Here, to investigate the involvement of the SPC in the cleavage event of Jaw1, Flp-In T-REx HEK293 cells lacking their catalytic subunits, SEC11A or SEC11C (hereafter called SEC11A KO and SEC11C KO cells, respectively), were produced by the CRISPR/Cas9 system (Fig. 4A,B). Single-guide RNAs (sgRNAs) were designed to target exon 2 of the respective genes, resulting in frameshift due to the indels upstream of the regions coding for catalytic residues. The loss of SEC11A was confirmed in two clones (#3 and #9) transfected with a plasmid coding for sgRNA against *seclla* by western blotting (Fig. 4C). On the other hand, no specific band corresponding to SEC11C was detected by western blotting, probably due to the low affinity (data not shown). Instead, by DNA sequencing, the homotypic 1-bp insertion was confirmed in two clones (#1 and #4) (Fig. 4D). In these cells, the FLAG Hu Jaw1 was expressed and the band patterns were compared. The result showed that the percentages of cleaved Jaw1 in SEC11A KO #3 and #9 cells were significantly reduced, whereas those of SEC11C KO #1 and #4 cells were comparable with wild-type cells (Fig. 4E,F). This result indicates that the C-terminal region of Jaw1 is cleaved by SPC with SEC11A but not SEC11C. To validate this, the rescue experiment was performed, in which it is tested whether or not the phenotypic changes due to the loss of genes are restored by re- or complementary protein expression. The expression of FLAG tagged SEC11A (FLAG SEC11A) in SEC11A KO #3 and #9 cells restored the percentage of cleaved Jaw1 compared with that of control cells, but that of FLAG SEC11C did not in spite of its much higher expression level compared to FLAG SEC11A (Fig. 4G–J). In summary, these data indicate that the SPC with SEC11A is specifically responsible for the cleavage event of the Jaw1 C-terminal region.

**Figure 4.**
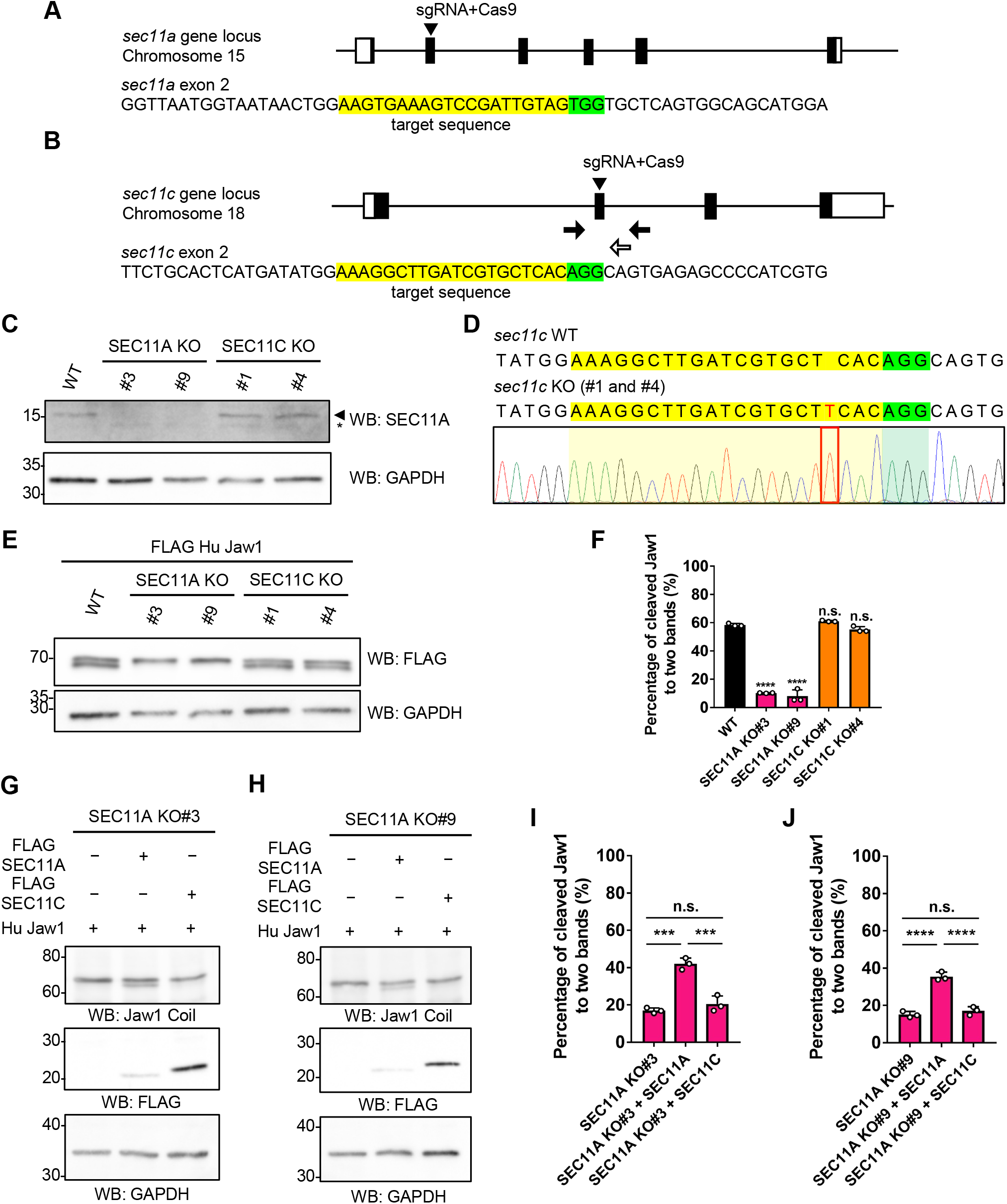
Identification of the protease responsible for Jaw1 C-terminal cleavage. **A, B**) Schematic diagrams indicating the generation of Flp-In T-REx HEK293 cells lacking *sec11a* (A) or *sec11c* (B) using the CRISPR/Cas9 system. The exons are represented as boxes (white, untranslated region; black, coding region). The arrowheads indicate the sgRNA targeting sites on the genomes. The arrows indicate the positions of the genotyping primers (closed triangles) and DNA sequencing (open triangles). **C**) Cell lysates of SEC11A KO #3 and #9 and SEC11C KO #1 and #4 cells were subjected to western blotting using an anti-SEC11A antibody. The arrowhead indicates the band of SEC11A. Asterisk; non-specific bands. **D**) The DNA sequence in the targeted site of *sec11c* of SEC11C KO cells. The indel is colored in red. **E**) FLAG Hu Jaw1 was expressed in SEC11A KO #3 and #9 and SEC11C KO #1 and #4 cells by transfection. After incubation for 24 h, the lysates were subjected to western blotting. **F**) Graph showing the percentage of Jaw1 C-terminal cleavage in (E). **G, H**) Hu Jaw1 was expressed alone or with either FLAG SEC11A or FLAG SEC11C in SEC11A KO #3 (G) and #9 (H) cells by transfection. After incubation for 24 h, the lysates were subjected to western blotting. **I, J**) Graph showing the percentage of Jaw1 C-terminal cleavage in (G) and (H), respectively. **F, I, J**) The averages of three independent experiments per condition are shown in the graphs. Error bar shows ±SD, “n.s.”, not significant; \*\*\**P* < 0.001; \*\*\*\**P* < 0.0001; statistical analysis, one-way ANOVA followed by Dunnett’s multiple comparison test (F) and Tukey’s multiple comparison test (I, J).

### Accessory subunits are involved in the Jaw1 C-terminal cleavage event

The SPC comprises three accessory subunits: SPCS1–3, in addition to a catalytic subunit (Liaci *et al*., 2021). To investigate the involvement of the SPC accessory subunits in the Jaw1 C-terminal cleavage event, Flp-In T-REx HEK293 Hu Jaw1 cells were treated with siRNAs against SPCS1–3 followed by doxycycline (Dox) addition to induce the expression of Jaw1. The cell lysates were then subjected to western blotting to evaluate the Jaw1 C-terminal cleavage event and the protein expression levels of SPC components. Prior to these investigations, the knockdown efficiency was measured by RT-qPCR. As shown in Figure 5A–C, the mRNA expression levels of *spcs1, spcs2*, and *spcs3* were significantly reduced to 20.9% in SPCS1 KD#1 cells, 2.0% in SPCS1 KD#2 cells, 50.0% in SPCS2 KD#1 cells, 6.0% in SPCS2 KD#2 cells, 5.1% in SPCS3 KD#1 cells and 5.4% in SPCS3 KD#2 cells, respectively, compared with control cells. Under this condition, the protein expression level of SPC components and the percentage of cleaved Jaw1 was evaluated. The protein expression levels of the SPC components: SPCS1, SPCS2, and SEC11A were also significantly reduced in SPCS1 KD#2 cells, SPCS2 KD #2 cells, and SPCS3 KD#1 and #2 cells (Fig. 5D–G). Furthermore, the percentage of cleaved Jaw1 was significantly reduced in SPCS1 KD#2 cells, SPCS2 KD#2 cells, SPCS3 KD#1 cells, and SPCS3 KD#2 cells (Fig. 5D,H). These results indicate that the depletion of SPC accessory subunits causes the reduction in the amount of other SPC components, and that the SPC accessory subunits are also involved in Jaw1 C-terminal cleavage.

**Figure 5.**
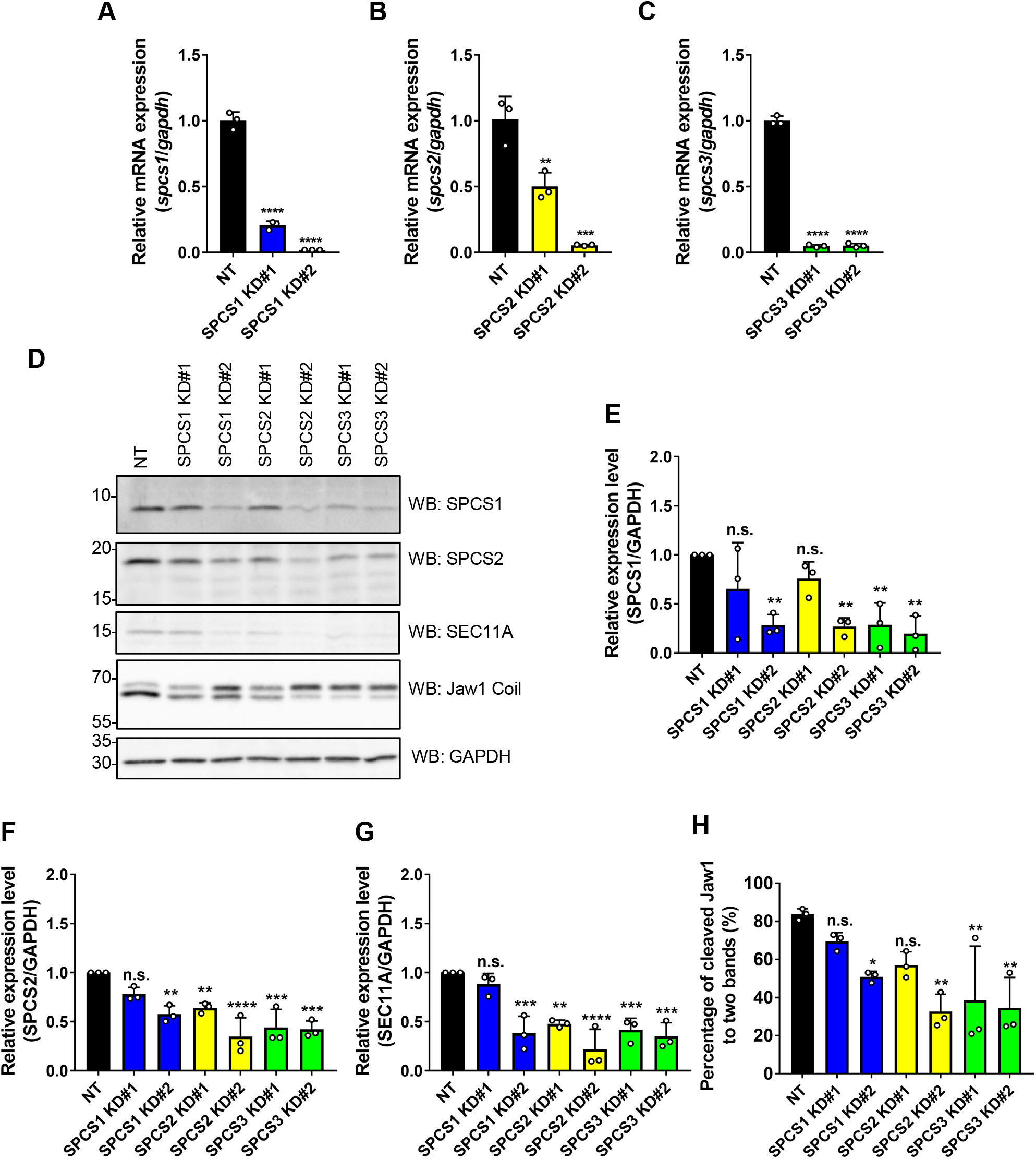
Investigation directed to the involvement of the SPC accessory subunits in Jaw1 C-terminal cleavage. Flp-In T-REx HEK293 Hu Jaw1 cells were treated with siRNA against *spcs1, spcs2*, and *spcs3* for 48 h. After 24 h from the start of siRNA treatment, the cells were treated with Dox for 24 h. **A-C**) Graphs showing the relative expression levels to NT of *spcs1* (A), *spcs2* (B), and *spcs3* (C) mRNA to *gapdh* measured by RT-qPCR. **D**) The cell lysates were subjected to western blotting. **E-G**) Graphs showing the relative expression level of SPCS1 (E), SPCS2 (F), and SEC11A (G) in (D). **H**) Graph showing the percentage of Jaw1 C-terminal cleavage in (D). **A-C, E-H**) The averages of three independent experiments per condition are shown in the graphs. Error bar shows ±SD, “n.s.”, not significant; \**P* < 0.05; \*\**P* < 0.01; \*\*\**P* < 0.001; \*\*\*\**P* < 0.0001. Statistics: one-way ANOVA followed by Dunnett’s multiple comparison test.

### The Jaw1 C-terminal region is cleaved after its insertion into the membrane

Next, we investigated whether the Jaw1 C-terminal region is cleaved post- or pre-insertion into the ER membrane. To evaluate this, we prepared the plasmids coding for FLAG Hu and Ms Jaw1 with a C-terminal opsin tag, which is glycosylated after its insertion into the ER membrane, as reported in previous studies (Fig. 6A) (Abell *et al*., 2007; Borgese *et al*., 2001; Coy-Vergara J *et al*., 2019; Favaloro *et al*., 2010; Masaki *et al*., 1996; Pedrazzini *et al*., 2000; Schuldiner *et al*., 2008; Stefanovic and Hegde, 2007). These proteins were expressed in HEK293 cells and the cell lysates were subjected to western blotting. Parts of the lysates were treated with EndoH prior to western blotting to characterize the bands corresponding to the glycosylated type; thereby, we could distinguish an uncleaved form of Jaw1 that was inserted into the ER membrane from the pre-inserted nascent polypeptide chain. As shown in Figure 6B and 6C, three bands were present in both lanes of FLAG Hu Jaw1 opsin and FLAG Ms Jaw1 opsin. Importantly, the upper bands, corresponding to the uncleaved Jaw1 with *N*-linked glycosylation disappeared after the treatment with EndoH. This result indicates that the C-terminal region of Jaw1 is cleaved after its insertion into the ER membrane. Furthermore, in both lanes of FLAG Hu AASS opsin and FLAG Ms AASS opsin, mutants with a deficit in the C-terminal cleavage event, the two bands were detected, corresponding to the upper and middle bands of FLAG Hu Jaw1 opsin and FLAG Ms Jaw1 opsin, thus, uncleaved Jaw1 with *N*-linked glycosylation and pre-inserted Jaw1, respectively (Fig. 6B,C). Thus, these results indicate that the C-terminal cleavage event is not required for the insertion into the ER membrane. In addition, the bands corresponding to uncleaved Jaw1 with *N*-linked glycosylation were detected in the SEC11A KO#3 and #9 cells and SPCS1-3 KD cells, although the ratios of cleavage event were changed due to the different depletion efficiency of the SPC component (Fig. 6D,E). In summary, the results suggest that the C-terminal region of Jaw1 is cleaved after its insertion into the ER membrane, and the cleavage event of the Jaw1 C-terminal region by the SPC is not involved in the insertion of Jaw1 into the ER membrane.

**Figure 6.**
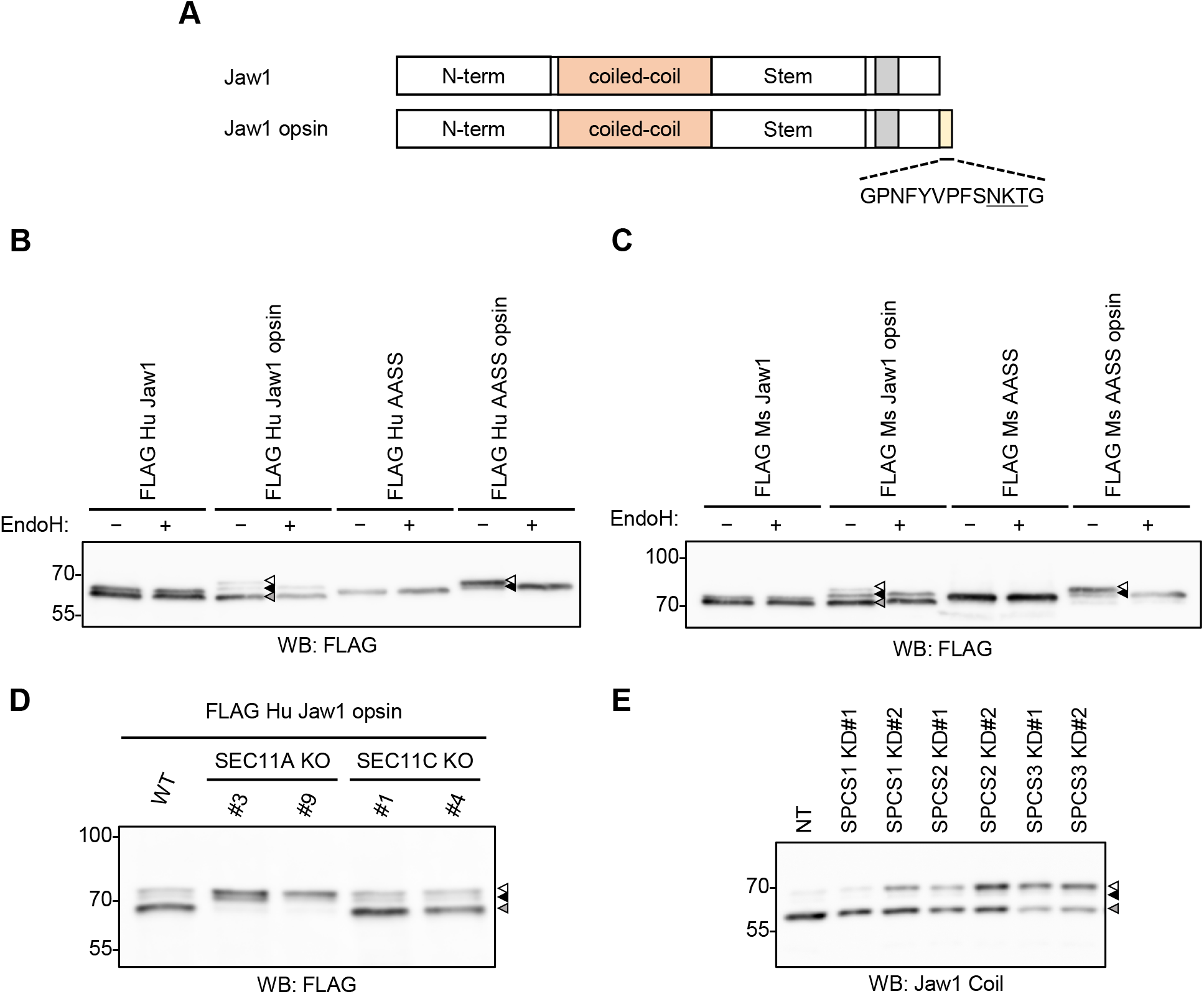
Investigation into the relevance of the Jaw1 C-terminal cleavage event to its traffic to the ER. **A**) Schematic representation of Jaw1 and Jaw1 opsin. The opsin tag was added to its C-terminal end. **B, C**) FLAG Hu Jaw1, FLAG Hu Jaw1 opsin, FLAG Hu AASS, and FLAG Hu AASS opsin (B) and FLAG Ms Jaw1, FLAG Ms Jaw1 opsin, FLAG Ms AASS, and FLAG Ms AASS opsin (C) were expressed in HEK293 cells by transfection. After incubation for 24 h, the lysates samples were treated with or without EndoH and subjected to western blotting using an anti-FLAG mouse antibody. **D**) FLAG Hu Jaw1 opsin was expressed in SEC11A KO #3 and #9 and SEC11C KO #1 and #4 cells by transfection. After incubation for 24 h, the lysates were subjected to western blotting using an anti-FLAG mouse antibody. **E**) Flp-In T-REx HEK293 Hu Jaw1 opsin cells were treated with siRNA against *spcs1, spcs2*, and *spcs3* for 48 h. After 24 h from the start of siRNA treatment, the cells were treated with Dox for 24 h. After that, the lysates were subjected to western blotting using an anti-Jaw1 Coil antibody. Hu Jaw1 opsin expressed by the treatment with Dox in this cell does not bear the N-terminal tags unlike FLAG Hu Jaw1 opsin in (B) and (D), thereby, an anti-Jaw1 Coil antibody but not an anti-FLAG mouse antibody was used. **B-E**) Open triangles, the bands of ER-inserted uncleaved Jaw1 with *N*-linked glycosylation; closed triangles, the bands of the pre-inserted Jaw1(black) and cleaved Jaw1 (gray). The representative blot images from three independent experiments with similar results are shown.

### The cleavage event of the Jaw1 C-terminal region enhances the augmentative effect of Jaw1 on the Ca^2+^ release from the ER

Finally, we investigated the significance of the Jaw1 C-terminal cleavage event. We previously reported that the C-terminal region of Jaw1 functions as a KASH domain to maintain the nuclear shape (Kozono *et al*., 2018). On the basis of the fact that the KASH domain is crucial for the interaction with SUN proteins, we hypothesized that the AASS mutant is strongly localized at the NE rather than the ER. To evaluate this, we prepared Jaw1 KO Flp-In T-REx HEK29 cells inducibly expressing Hu Jaw1 or Hu AASS (hereafter called Jaw1 IE or AASS IE) followed by immunostaining. The confocal images showed that the AASS mutant was localized at both the NE and the ER, similar to wild-type Jaw1 (Fig. 7A). The nuclear shape in AASS IE cells was also the same as in Jaw1 IE cells, indicating that the AASS mutant had no dominant negative effect on the nuclear shape. IP_3_Rs release Ca^2+^ from the ER into the cytoplasm when the IP_3_ is produced upon GPCR stimulation (Fig. S3). Jaw1 has an additional role to increase the Ca^2+^ release from the ER via interaction with IP_3_Rs upon GPCR stimulation, as we recently reported (Okumura *et al*., 2022). Therefore, we measured Ca^2+^ flux in Jaw1 KO Flp-In T-REx HEK293 (Jaw1 KO), Jaw1 IE, and AASS IE cells via stimulation with ATP. Prior to the measurement, it was confirmed that the protein expression levels of IP_3_R1-3 and Jaw1 were not markedly changed among cell lines (Fig. 7B–F). Furthermore, it was confirmed by the co-immunoprecipitation assay that the interaction with IP_3_Rs was maintained in both Hu Jaw1 and AASS mutant cells (Fig. S4). In this context, Ca^2+^ imaging was performed. As shown in Figure 7G, the kinetic curves of Ca^2+^ flux maintained higher values in both Jaw1 IE and AASS IE cells compared with Jaw1 KO cells in response to stimulation with 100 μM ATP solution. Furthermore, the maximum amplitude and area under the curve (AUC) significantly increased in both Jaw1 IE cells and AASS IE cells compared with Jaw1 KO cells (Fig. 7H,I). However, the kinetic curves of Ca^2+^ flux in AASS IE cells maintained a slightly lower value than that of Jaw1 IE cells (Fig. 7G). In addition, the maximum amplitude and AUC of AASS IE cells were significantly lower than that of Jaw1 IE cells (Fig. 7H,I). These phenotypic differences in Ca^2+^ flux between Jaw1 IE cells and AASS IE cells were observed in response to stimulation with a lower concentration of ATP (Fig. S5). These results indicate that the ability to increase the Ca^2+^ release from the ER in the AASS mutant was slightly lower than that of wild-type Jaw1. In other words, the cleavage event of the Jaw1 C-terminal region enhances the augmentative effect of Jaw1 on the Ca^2+^ release from the ER via the interaction with IP_3_Rs.

**Figure 7.**
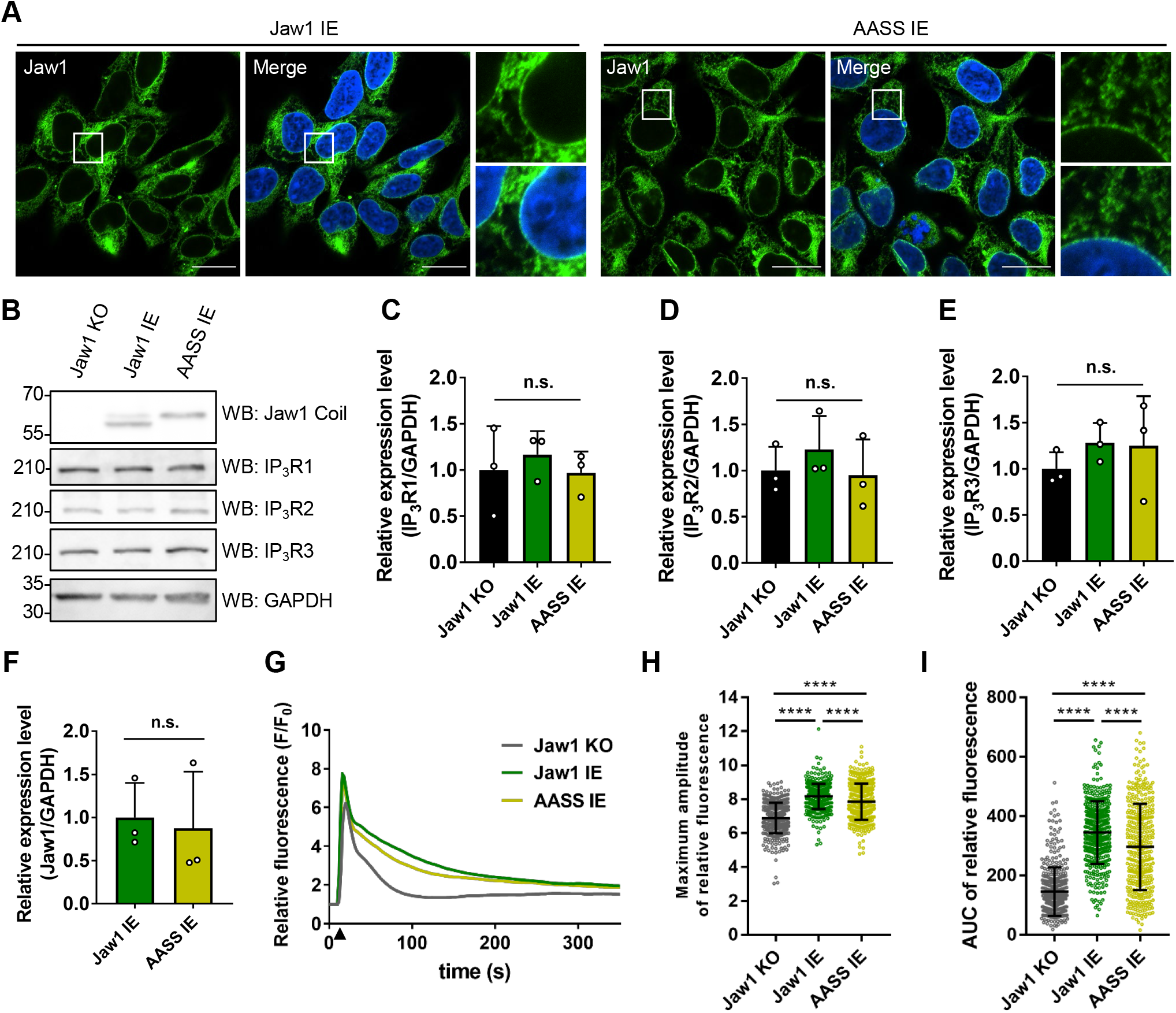
Investigation into the relevance of Jaw1 C-terminal cleavage event in Ca^2+^ release via IP_3_Rs. Jaw1 KO, Jaw1 IE, and AASS IE cells were treated with Dox for 24 h. **A**) The cells were used for immunostaining using an anti-Jaw1 N antibody (green). Nuclei were stained with Hoechst33342 (blue). The images were acquired by confocal microscopy. Scale bars: 20 μm. The magnified images corresponding to the area surrounded with white lines in each image were generated. **B**) The lysates were subjected to western blotting. **C-F**) Graphs showing the relative expression level of IP_3_R1 (C), IP_3_R2 (D), IP_3_R3 (E), and Jaw1 (F) in (B). The averages of three independent experiments per condition are shown. **G-I**) The cells were subjected to Ca^2+^ imaging. **G**) Mean curves of relative Fluo-4 intensity. Closed triangles represent the time points that 100 μM ATP solution was added. **H, I**) The maximum amplitude (H) and AUC (I) of relative Fluo-4 intensity (n = 400). **C-F, H, I**) Error bar shows ±SD, “n.s.”, not significant; \*\*\*\**P* < 0.0001. Statistics: two-tailed Student’s *t*-test (F) and one-way ANOVA followed by Tukey’s multiple comparison test (C–E, H, I).

## Discussion

In this study, we identified two alanine residues (509/510) as a cleavage site of the Jaw1 C-terminal region by mutational analysis and protein sequencing. Furthermore, using cells that lacked SPC components, we showed that the SPC with SEC11A but not SEC11C specifically cleaves Jaw1. Importantly, the analysis using opsin-tagged mutants indicated that the cleavage event occurs after its insertion into the ER membrane and that the cleavage event is not required for the traffic and insertion of Jaw1 into the ER membrane. Finally, our data in the functional assay demonstrated that the augmentative effect of Jaw1 on Ca^2+^ release from the ER is slightly lower in cells that express the AASS mutant, and did not affect the localization of the NE and the ER similar to that of wild-type Jaw1. In summary, the results suggested that the cleavage event of the Jaw1 C-terminal region by the SPC enhances its ability to increase the Ca^2+^ release from the ER via interaction with IP_3_Rs.

The results of the mutational analysis and protein sequencing identified the cleavage site of Jaw1 (…TAVEA/APTQEGD…) (Fig. 3). Signal peptides typically contain three distinctive regions: the *n*-region, which carries a positive charge; the *h*-region, which comprises abundant hydrophobic residues; and the *c*-region, which contains polar residues. The signal peptides processed by the SPC contain small uncharged residues at positions −1 and −3 to the cleavage site in the *c*-region (Paetzel *et al*., 2002), and the identified cleavage site of Jaw1 (… TA<u>V</u>E<u>A</u>/APTQEGD…) is also consistent with this rule. Interestingly, a second peptide (AP/TQEGD…) was also detected by protein sequencing. We speculated that its presence was due to further processing by other enzymes. It has been reported that the cleaved C-terminal peptide is delivered to major histocompatibility complex class I (Snyder *et al*., 1997). The further processing by other enzymes following the processing by SPC might be crucial in transforming the C-terminal peptide into a bioactive type for its function.

It is canonically known that the SPC cleaves off the amino-terminal signal peptides dependent on signal recognition particle and SEC61 (Paetzel *et al*., 2002). Although it is known that the C-terminal region of synaptobrevin, one of the TA proteins, is cleaved by SPC (Nilsson *et al*., 2002), how TA proteins are processed by the SPC remains unclear. In this context, Jaw1 would be a new model protein to investigate the cleavage event of TA proteins by SPC. Furthermore, our results showed that SPCs with SEC11A but not SEC11C specifically cleaved the C-terminal region of Jaw1 (Fig. 4). To the best of our knowledge, this is the first study reporting the differences in the substrate specificity of SEC11A and SEC11C. The sequential identity between them is high (79.3%), and the amino acids in the transmembrane domain and the catalytic residues are completely conserved; moreover, the stability of SPC with SEC11C is slightly lower than that with SEC11A (Liaci *et al*., 2021). The subtle differences in amino acid composition and stability might affect their catalytic specificity, perhaps especially in the case of the TA proteins. In addition, the depletion of the accessory subunit resulted in the reduction in the percentage of the Jaw1 C-terminal cleavage (Fig. 5). Behind this, other SPC components were also reduced, indicating that the accessory subunits of mammalian SPCs have a role in maintaining the complex’s stability, consistent with the previous report on yeast SPCs (Meyer and Hartmann, 1997).

Although six KASH proteins, including Jaw1, are known (Kozono *et al*., 2018; Morimoto *et al*., 2012; Starr, 2009), the cleavage event of the C-terminal region is unique among them (Fig. 1). This result led us to a new hypothesis that the cleavage event generates the uncleaved form of Jaw1 with the KASH domain and the cleaved form of Jaw1 without the KASH domain, thereby resulting in the distinctive localization at the NE and ER, and that Jaw1 functions as a KASH protein and an IP_3_R regulator. However, the AASS mutant was localized at both the ER and NE, comparable with the localization of wild-type Jaw1 (Fig. 7A). This might mean that the affinity of the Jaw1 KASH domain to interact with SUN proteins is weaker than that of other KASH proteins, although all KASH domains have an affinity for SUN proteins (Cruz *et al*., 2020). Thus, the preferential interaction of SUN proteins with other KASH proteins might result in the location of uncleaved form Jaw1 to the ER as well as NE. Furthermore, the intracellular extent of KASH proteins and SUN proteins would be an important factor affecting the localization of Jaw1. Intriguingly, their expression level is heterogeneous depending on the tissue type and cell differentiation stage (Göb *et al*., 2010; Morimoto *et al*., 2012; Olins *et al*., 2009; Rajgor *et al*., 2012; Yang *et al*., 2018). In addition, Jaw1 is specifically expressed in taste cells on the tongue, small intestinal tuft cells, pancreatic acinar cells, and immune cells (Behrens *et al*., 1994; Chang *et al*., 2021; Prüschenk *et al*., 2021; Shindo *et al*., 2010). Further analysis using the cell types expressing Jaw1 endogenously would provide deeper insight into the relationship between the cleavage event of the Jaw1 C-terminal region and its localization.

We previously reported that Jaw1 interacts with IP_3_Rs via its coiled-coil domain, and a mutant lacking the coiled-coil domain, has no augmentative effect of Jaw1 on the Ca^2+^ signaling, which indicates that Jaw1 directly increases the Ca^2+^ release activity of IP_3_Rs (Okumura *et al*., 2022). In this study, the augmentative effect of the AASS mutation on the Ca^2+^ release from the ER was slightly lower than that of wild-type Jaw1 (Fig. 7). Based on these data, we suggest that the C-terminal cleavage event of Jaw1 is crucial step for its maturation to function as an IP_3_R regulator, although we were not able to elucidate the mechanism. Considering the unchanged localization and maintained interaction with the IP_3_Rs of the AASS mutant compared with the wild-type, the reduction in Ca^2+^ release from the ER in AASS IE cells may be due to structural hindrance in the Ca^2+^ pathway of the IP_3_R channels by the C-terminal region of Jaw1. To unveil this, structural approaches such as cryo-transmission electron microscopy would be required. On the other hand, we could not completely exclude the possibility that Jaw1 augments the Ca^2+^ signaling by affecting other proteins such as Ca^2+^ pumps, transporters, and Ca^2+^ buffering in addition to the activity of IP_3_Rs, and the slightly lower effect of the AASS mutant on the Ca^2+^ signaling is brought by unknown factors except IP_3_Rs. The calcium signaling in the taste cells, small intestinal cells, and pancreatic acinar cells, where Jaw1 is specifically expressed, is essential for their physiological functions, such as taste perception, helminth expulsion, and exocrine secretion (Futatsugi *et al*., 2005; Hisatsune *et al*., 2007; Luo *et al*., 2019; McGinty *et al*., 2020; Nadjsombati *et al*., 2018). It would be also required to investigate whether the cleavage event of Jaw1 might significantly affect the Ca^2+^ release from the ER in these cells and their physiological function.

## Materials and Methods

### Plasmids

The plasmids, pcDNA5 FRT/TO HA FLAG Ms Jaw1 (Kozono *et al*., 2018), pcDNA5 FRT/TO HA FLAG Hu Jaw1 (Kozono *et al*., 2021), pcDNA3.1 (+) Hu Jaw1 (Kozono *et al*., 2021), and pcDNA5 FRT/TO Hu Jaw1 (Okumura *et al*., 2022), were produced as described in previous studies. The primer sets used to produce the plasmids are shown in Table S1. For the generation of the plasmids encoding FLAG Ms Jaw1 KASH chimeras, the DNA fragments coding for each luminal region of the KASH protein were first amplified by PCR from a total cDNA library derived from mouse (C57BL/6J) kidney (K1 and K2), using primer sets 1 and 2, and from plasmids (pcDNA5 FRT/TO HA FLAG Ms Nesprin3 (K3), Nesprin4 (K4) and KASH5 (K5)) using primer sets 3–5, respectively. The PCR products were subcloned into the pcDNA5 FRT/TO HA FLAG Ms Jaw1 (digested with *Afe*I/*Bam*HI) using an In-Fusion HD Cloning Kit (#Z9648N; TaKaRa, Kusatsu, Japan) to swap the luminal region of Jaw1 to that of KASH proteins, resulting in pcDNA5 FRT/TO HA FLAG Ms Jaw1 K1-K5. For the production of the plasmids encoding the mutants in which the specific five residues in the luminal region of Jaw1 were substituted with alanine residues, the DNA fragments were first amplified by PCR with primer sets 6/7, 8/9, 10/11, 12/13, 14/15, 16/17, 18/19, and 20/21 from pcDNA5 FRT/TO HA FLAG Ms Jaw1. The PCR products were subcloned into the pcDNA5 FRT/TO HA FLAG Ms Jaw1 (digested with *Hpa*I/*Sph*I) by double-insert cloning using an In-Fusion HD Cloning Kit (#Z9648N; TaKaRa, Kusatsu, Japan), resulting in pcDNA5 FRT/TO HA FLAG Ms Jaw1 504–508A, 507–511A, 512–516A, 517–521A, 522–526A, 527–531A, 532–536A, and 535–539A, respectively. For the production of pcDNA5 FRT/TO HA FLAG Ms AASS and pcDNA5 FRT/TO HA FLAG Hu AASS, the DNA fragments were first amplified by PCR with primer sets 22/23 and 24/25 from pcDNA5 FRT/TO HA FLAG Ms Jaw1 and pcDNA5 FRT/TO HA FLAG Hu Jaw1. The PCR products were subcloned into the pcDNA5 FRT/TO HA FLAG Ms Jaw1 (digested with *Hpa*I/*Sph*I) and pcDNA5 FRT/TO HA FLAG Hu Jaw1 (digested with *Hpa*I/*Sph*I), respectively, by double-insert cloning using an In-Fusion HD Cloning Kit (#Z9648N; TaKaRa, Kusatsu, Japan). The pcDNA5 FRT/TO HA FLAG Hu AASS was then digested with *Bam*HI/*Apa*I and the DNA fragment was ligated into the pcDNA5 FRT/TO vector using a DNA Ligation Kit (#6023; TaKaRa, Kusatsu, Japan), resulting in pcDNA5 FRT/TO Hu AASS. For the generation of pcDNA5 FRT/TO HA FLAG Ms Jaw1 opsin and pcDNA5 FRT/TO HA FLAG Ms AASS opsin, the DNA fragments coding for C-terminal opsin-tagged Ms Jaw1 were first amplified by PCR with primer set 26 from pcDNA5 FRT/TO HA FLAG Ms Jaw1 and pcDNA5 FRT/TO HA FLAG Ms AASS. The PCR products were subcloned into the pcDNA5 FRT/TO HA FLAG Ms Jaw1 (digested with *Hpa*I/*Bam*HI) using an In-Fusion HD Cloning Kit (#Z9648N; TaKaRa, Kusatsu, Japan). Similarly, the DNA fragments coding for C-terminal opsin-tagged Hu Jaw1 were amplified by PCR with primer set 27 from pcDNA5 FRT/TO HA FLAG Hu Jaw1 and pcDNA5 FRT/TO HA FLAG Hu AASS. The PCR products were subcloned into pcDNA5 FRT/TO HA FLAG Hu Jaw1 or pcDNA5 FRT/TO Hu Jaw1 (digested with *Hpa*I/*Xho*I) using an In-Fusion HD Cloning Kit (#Z9648N; TaKaRa, Kusatsu, Japan), resulting in pcDNA5 FRT/TO HA FLAG Hu Jaw1 opsin, pcDNA5 FRT/TO HA FLAG Hu AASS opsin, and pcDNA5 FRT/TO Hu Jaw1 opsin. To generate the plasmids coding for PA-tagged Ms Jaw1, the pcDNA3.1 (+) C-terminal PA vector was first prepared by insertion of a DNA cassette encoding the PA tag (annealed primer set 28) into the pcDNA3.1(+) vector (digested with *Xho*I/*Xba*I) using a DNA Ligation Kit (#6023; TaKaRa, Kusatsu, Japan). The DNA fragment coding for Ms Jaw1 was then amplified by PCR with primer set 29 from pTagGFP2-C Ms Jaw1 (previously described by Kozono *et al*., 2018) and the PCR product was subcloned into the pcDNA3.1 (+) C-terminal PA vector (digested with *Kpn*I/*Xho*I) using an In-Fusion HD Cloning Kit (#Z9648N; TaKaRa, Kusatsu, Japan), resulting in pcDNA3.1(+) Ms Jaw1 PA. Furthermore, the DNA fragment coding for the N-terminal region of Ms Jaw1 was amplified by PCR with primer set 30 from pcDNA5 FRT/TO HA FLAG Ms Jaw1, and the PCR product was subcloned into the pcDNA3.1(+) Ms Jaw1 PA (digested with *Xho*I) using an In-Fusion HD Cloning Kit (#Z9648N; TaKaRa, Kusatsu, Japan), resulting in pcDNA3.1(+) Ms Jaw1 NIDR PA. Finally, pcDNA3.1(+) Ms Jaw1 PA and pcDNA3.1(+) Ms Jaw1 NIDR PA were digested with *Kpn*I/*Apa*I, and the DNA fragments were ligated into the pcDNA5 FRT/TO HA FLAG vector (digested with same enzymes) using a DNA Ligation Kit (#6023; TaKaRa, Kusatsu, Japan), resulting in pcDNA5 FRT/TO HA FLAG Ms Jaw1 PA and pcDNA5 FRT/TO HA FLAG Ms Jaw1 NIDR PA, respectively. For the production of pSpCas9-2A-Puro sec11a or sec11c sgRNA, the DNA cassettes (annealed primer set 31 and 32, respectively) containing the sgRNA sequence targeting the coding region of the *sec11a* and *sec11c* genes were inserted into the pSpCas9(BB)-2A-Puro (PX459) vector (#48139; Addgene, Watertown, MA, USA) (digested with *Bbs*I) using a DNA Ligation Kit (#6023; TaKaRa, Kusatsu, Japan). For the cloning of the *sec11a* (NM_014300.3) and *sec11c* (NM_033280.3) genes, the DNA fragment coding for an entire sequence of *sec11a* and *sec11c* genes were amplified by PCR with primer sets 33 and 34 from a total cDNA library derived from HEK293 cells, and the PCR products were subcloned into the pcDNA3.1(+) vector (digested with *Kpn*I/*Bam*HI) using an In-Fusion HD Cloning Kit (#Z9648N; TaKaRa, Kusatsu, Japan), resulting in pcDNA3.1(+) Hu *sec11a* and pcDNA3.1(+) Hu sec11c, respectively. These plasmids were digested with *Kpn*I/*Bam*HI, and the DNA fragments were ligated to the pcDNA5 FRT/TO HA FLAG vector (digested with the same enzymes) using a DNA Ligation Kit (#6023; TaKaRa, Kusatsu, Japan), resulting in pcDNA5 FRT/TO HA FLAG Hu sec11a and pcDNA5 FRT/TO HA FLAG Hu sec11c. The identity of the cloned sequences was verified by a DNA sequence contract service (Genewiz, Chelmsford, MA, USA). After the confirmation, each plasmid was prepared using a Plasmid Mini Kit (#12123; QIAGEN, Germantown, MD, USA) and used for the subsequent experiments.

### Establishment of cell lines

Jaw1 KO Flp-In T-REx HEK293 (Jaw1 KO) cells and Jaw1 KO Flp-In T-REx HEK293 Hu Jaw1 (Jaw1 IE) cells was produced as previously described (Okumura *et al*., 2022). Flp-In T-REx HEK293 FLAG Ms Jaw1, FLAG Ms Jaw1 K1-K5, Hu Jaw1, Hu Jaw1 opsin, and Hu AASS were produced by co-transfection of pOG44 Flp-Recombinase Expression Vector (#V600520; Thermo Fisher Scientific, Waltham, MA, USA) with pcDNA5 FRT/TO HA FLAG Ms Jaw1, pcDNA5 FRT/TO HA FLAG Ms Jaw1 K1-K5, pcDNA5 FRT/TO Hu Jaw1, pcDNA5 FRT/TO Hu Jaw1 opsin and pcDNA5 FRT/TO Hu AASS, respectively, into the Flp-In™ T-REx™ HEK293 cell line or Jaw1 KO cells (#R78007; Thermo Fisher Scientific, Waltham, MA, USA). After incubation for 24 h, the medium was refreshed. The cells were then subjected to selection by treatment with 100 μg ml^−1^ of hygromycin B (#07296-66; Nacalai Tesque, Kyoto, Japan) for 48 h. After the medium was refreshed again, the cells were expanded and used for the subsequent assay. For the inducible gene expressions, the cells were treated with 200 ng ml^−1^ of Dox (#631311; TaKaRa, Kusatsu, Japan) for 24 h. For the generation of Flp-In T-REx HEK293 lacking *sec11a* or *sec11c*, Flp-In T-REx HEK293 cells were transfected with pSpCas9-2A-Puro sec11a or sec11c sgRNA. After incubation for 24 h, the medium was refreshed, and the cells were subjected to selection by treatment with 4 μg ml^−1^ of puromycin (#P8833; Sigma-Aldrich, St Louis, USA). The cells were then single colonized by the limiting dilution cloning method in 96-well plates and expanded. For the verification of the mutation by Sanger sequencing, the genomes were purified from the cells using a QIAamp DNA Micro Kit (#51304; QIAGEN, Germantown, MD, USA), and the amplified PCR products containing the target region using primer set 35 were submitted to the DNA sequencing contract service (Genewiz, Chelmsford, MA, USA).

### Cell culture

HEK293 cells and the established Flp-In T-REx HEK293 cells were cultured in DMEM (#05919; Nissui Pharmaceuticals Co. Ltd., Tokyo, Japan) containing 10% fetal bovine serum, 100 U ml^−1^ penicillin, 100 μg ml^−1^ streptomycin (#168-23191; FUJIFILM Wako Pure Chemical Corporation, Tokyo, Japan), and 5.84 mg ml^−1^ _L_-glutamine (#16919-42; Nacalai Tesque, Kyoto, Japan) at 37°C and 5% CO_2_.

### Transfection

Plasmids were introduced into the cells using Screen*F*ect™A *plus* (#299-77103; FUJIFILM Wako Pure Chemical Corporation, Tokyo, Japan) according to the manufacturer’s instructions. After the treatment for 24 h, the cells were used for the subsequent assays.

### RNA interference

siRNA-mediated knockdown assays were performed using Lipofectamine™ RNAiMAX transfection reagent (#13778030; Thermo Fisher Scientific, Waltham, MA, USA) according to the manufacturer’s instructions. Custom oligonucleotides specific for human *spcs1* #1 (5’-UGGUUUUAGCUCUUAUCUGGG-3’, 5’-CAGAUAAGAGCUAAAACCACC-3’), human *spcs1* #2 (5’-GUUAUGGCCGGAUUUGCUUtt-3’, 5’-AAGCAAAUCCGGCCAUAACtc-3’), human *spcs2* #1 (5’-UUACUUUACCCAUUUUAAGUU-3’, 5’-CUUAAAAUGGGUAAAGUAAGA-3’), human *spcs2* #2 (5’-UAGGAUAUGACACACAAAGCC-3’, 5’-CUUUGUGUGUCAUAUCCUAUU-3’), human *spcs3* #1 (5’-UAUCAAAUGUGAUAAAUCCCA-3’, 5’-GGAUUUAUCACAUUUGAUAUA-3’), and human *spcs3* #2 (5’-UCGUUAUUUCAUAUGUAUCUG-3’, 5’-GAUACAUAUGAAAUAACGAAG-3’) were used. For the control of RNA interference, stealth control 172 (Invitrogen, Carlsbad, CA, USA) was used. After the treatment with siRNA for 48 h, the cells were subjected to the subsequent assays.

### RT-qPCR

RT-qPCR was performed as previously described with some modifications (Kozono *et al*., 2018). In brief, the cells were grown in a 6-well plate (#TR5000; NIPPON Genetics Co. Ltd., Tokyo, Japan), and total RNA was isolated using RNAiso Plus (#9108; TaKaRa, Kusatsu, Japan) according to the manufacturer’s instructions. Total RNA (400 ng) was then reverse-transcribed using PrimeScript RT Master Mix (#RR036A; TaKaRa, Kusatsu, Japan). The cDNA was then subjected to the subsequent PCR using TB Green *Premix Ex*™ Taq II (Tli RNaseH Plus) (#RR820A; TaKaRa, Kusatsu, Japan) on a Thermal Cycler Dice Real Time System II MRQ (TP960; TaKaRa, Kusatsu, Japan) or Thermal Cycler Dice Real Time System (TP800; TaKaRa, Kusatsu, Japan), using the following primers: *spcs1* _forward; 5’-ATCTACGGGTACGTGGCTGA-3’, *spcs1*_reverse; 5’-GCGATAGATGGGCCATGGAG-3’, *spcs2*_forward; 5’-TGACAAGTGGGATGGATCAGC-3’, *spcs2*_reverse; 5’-TACAGATGGTGAGGCGACCA-3’, *spcs3*_forward; 5’-CTCACTGTTCGCCTTCTCGC-3’, *spcs3*_reverse; 5’-TCCTGTCTTTGAAGGCGGTG-3’, *gapdh*_forward; 5’-GAGTCCACTGGCGTCTTCAC-3’, and *gapdh*_reverse; 5’-GGTGCTAAGCAGTTGGTGGT-3’.

### Immunostaining

Immunostaining was performed as previously described with some modifications (Kozono *et al*., 2021). In brief, the cells were grown on a glass-bottom dish (#D141400; MATSUNAMI, Osaka, Japan), washed with PBS once, and fixed in 4% paraformaldehyde/PBS for 10 min. After washing with PBS, the cells were permeabilized using 0.1% Triton X-100/PBS for 5 min and blocked with 3% bovine serum albumin (BSA)/PBS for 1 h. The cells were then reacted with the following primary antibodies diluted to 1:400 with 1% BSA/PBS for 1 h: anti-Jaw1 N rabbit antibody (produced in our laboratory as described (Okumura *et al*., 2022)) and anti-FLAG mouse antibody (#014-23383; FUJIFILM Wako Pure Chemical Corporation, Tokyo, Japan). After washing with 0.1% BSA/PBS three times for 5 min, the cells were incubated with the following secondary antibodies diluted to 1:500 with 0.1% BSA/PBS containing Hoechst33342 for 1 h: Alexa Fluor™ 488 goat anti-rabbit IgG (H+L) (#A11008; Thermo Fisher Scientific, Waltham, MA, USA) and Alexa Fluor™ 488 goat anti-mouse IgG (H+L) (#A11001; Thermo Fisher Scientific, Waltham, MA, USA). After washing with 0.1% BSA/PBS three times for 5 min, the cells were immersed in 0.1% BSA/PBS. The confocal images were acquired using a confocal microscope (LSM710; Carl Zeiss, Oberkochen, Germany) (objective lens; Zeiss Plan Apo-chromat 63 × 1.4 NA) or (AXR; Nikon, Tokyo, Japan) (objective lens; Plan Apo l 60 × 1.40 oil).

### Western blotting

Western blotting was performed as previously described with some modifications (Kozono *et al*., 2021). In brief, the cells were grown in a 6-well plate (#TR5000; NIPPON Genetics Co. Ltd., Tokyo, Japan), peeled off by gentle pipetting in 1 ml PBS, and centrifuged at 500 × *g* for 5 min at 4°C. The pellets were lysed in lysis buffer (50 mM Tris–HCl pH 7.6, 150 mM NaCl, 1% NP-40) containing 1 μL of a protease inhibitor cocktail (#25955-24; Nacalai Tesque, Kyoto, Japan). The lysates were sonicated on ice for 10 min, then centrifuged at 12,000 × *g* for 20 min at 4°C. The supernatants were mixed with SDS-PAGE buffer, and heat blocked at 95°C for 5 min. For the confirmation of glycosylation in the opsin-tagged mutants, the samples were mixed with Endo Hf (#P0703; New England Biolabs, Beverly, MA, USA), incubated at 37°C for 1 h, and heat blocked at 95°C for 5 min. The samples were subjected to SDS-PAGE followed by blotting on an Amersham™ Hybond™ P 0.45-μm PVDF membrane (#10600029; Cytiva, Marlborough, MA, USA). For the detection of SEC11A, SPCS1, and SPCS2, the samples were blotted onto a Amersham™ Hybond™ P 0.2-μm PVDF membrane (#10600058; Cytiva, Marlborough, MA, USA). The membranes were blocked in 3% skim milk (#190-12865; FUJIFILM Wako Pure Chemical Corporation, Tokyo, Japan) diluted with Tris-buffered saline (TBS) (20 mM Tris–HCl pH 7.6 and 137 mM NaCl) containing 0.1% Tween-20 (TBS-T) for 1 h. After washing the membrane with TBS-T, it was reacted with the following primary antibodies diluted with 1% skim milk/TBS-T overnight at 4°C: anti-FLAG mouse antibody (1:1000) (#014-23383; FUJIFILM Wako Pure Chemical Corporation, Tokyo, Japan), anti-GAPDH mouse monoclonal antibody (1:1000) (#016-25523; FUJIFILM Wako Pure Chemical Corporation, Tokyo, Japan), anti-Jaw1 Coil rat antibody (1:500 or 1:1000) (produced in our laboratory as previously described (Kozono *et al*., 2018)), anti-SEC11A rabbit polyclonal antibody (1:1000) (#14753-1-AP; Proteintech, Wuhan, Hubei, China), anti-SPCS1 rabbit polyclonal antibody (1:1000) (#11847-1-AP; Proteintech, Wuhan, Hubei, China), anti-SPCS2 rabbit polyclonal antibody (1:1000) (#14872-1-AP; Proteintech, Wuhan, Hubei, China), anti-Jaw1 N rabbit antibody (1:1000) (produced in our laboratory as described (Okumura *et al*., 2022)), anti-IP_3_R1 rabbit antibody (1:1000) (#A7905; ABclonal, Wuhan, China), anti-IP_3_R2 mouse antibody (1:1000) (#sc-398434; Santa Cruz Biotechnology, CA, USA), and anti-IP_3_R3 mouse antibody (1:1000) (#610312;BD Bioscience, CA, USA). After washing the membrane with TBS-T, it was incubated with secondary antibodies diluted into 1:5000 with TBS-T for 1 h: Anti-Rat IgG, HRP-Linked Whole Ab Goat (#NA935; Cytiva, Marlborough, MA, USA), Anti-Rabbit IgG, HRP-Linked Whole Ab Donkey (#NA934; Cytiva, Marlborough, MA, USA), and anti-mouse IgG, HRP-Linked Whole Ab Sheep (#NA931; Cytiva, Marlborough, MA, USA). The membrane was then washed with TBS-T and reacted with ImmunoStar Zeta (#295-72404; FUJIFILM Wako Pure Chemical Corporation, Tokyo, Japan) or SuperSignal™ West Atto Ultimate Sensitivity Chemiluminescent Substrate (#A38555; Thermo Fisher Scientific, Waltham, MA, USA). The bands were detected using the LAS4000 Imaging System (GE Healthcare, Freiburg, Germany) or iBright Imaging System (Thermo Fisher Scientific, Waltham, MA, USA). The band intensities were calculated from the acquired images using Fiji (https://imagej.net/Fiji). The cropped blot images in each figure were generated from the corresponding full-length blots, as shown in Blot transparency (Fig. S6–S12).

### Immunoprecipitation

Co-immunoprecipitation assay to investigate the interaction of Jaw1 and IP_3_Rs was performed as previously described with some modifications (Okumura *et al*., 2022). In brief, the cells were grown in a 100-mm dish (#TR4002; NIPPON Genetics Co. Ltd., Tokyo, Japan), peeled off by gentle pipetting in 1 ml of PBS, and centrifuged at 500 × *g* for 5 min at 4°C. The pellets were lysed in lysis buffer (50 mM Tris–HCl pH 7.6, 150 mM NaCl, 0.5% NP-40) containing 1 μL of a protease inhibitor cocktail (#25955-24; Nacalai Tesque, Kyoto, Japan). The lysates were sonicated on ice for 10 min and centrifuged at 12,000 × *g* for 20 min at 4°C. A portion of the supernatant was mixed with SDS-PAGE buffer and heat blocked at 95°C for 5 min, resulting in the input samples. The remaining supernatant was diluted with lysis buffer containing 0.1% NP-40 and then incubated with anti-Jaw1 N antibody-conjugated beads by rotation for 1 h at 4°C. The conjugated beads were prepared by mixing anti-Jaw1 N antibody and Protein A Mag Sepharose™ (#28944006; Cytiva, Marlborough, MA, USA) for 1 h at 4°C. The beads that reacted with cell lysate were then collected by a magnetic stand and washed with wash buffer (50 mM Tris–HCl pH 7.6, 150 mM NaCl) three times. The beads were then mixed with SDS-PAGE buffer and heated at 95°C for 5 min. The supernatants were collected on the magnetic stand as immunoprecipitated samples and were subjected to western blotting as described above. For the immunoprecipitation of PA-tagged proteins, the cells were cultured on 100-mm dishes (25 dishes), and the PA-tagged proteins were expressed by transfection. After incubation for 24 h, the lysates were prepared as described above. The lysates were incubated with Anti-PA-tag Antibody Beads (#012-25841; FUJIFILM Wako Pure Chemical Corporation, Tokyo, Japan) by rotation for 1 h at 4°C. The beads were then collected by brief centrifugation and washed with wash buffer four times. For elution, the beads were mixed with SDS-PAGE buffer and heated at 95°C for 5 min. After brief centrifugation, the supernatants were collected as immunoprecipitated samples.

### Protein sequencing

Immunoprecipitated materials with anti-PA-tag antibody were separated by SDS-PAGE, electroblotted onto an Amersham™ Hybond™ P 0.45-μm PVDF membrane (#10600029; Cytiva, Marlborough, MA, USA), and stained with CBB R-250. The protein band corresponding to Fragment 2 was cut out and subjected to N-terminal sequencing analysis (Matsudaira, 1987) on an automated protein sequencer (#Procise 491 cLc; Applied Biosystems, Foster City, CA, USA) using the standard pulsed-liquid program for PVDF-blotted proteins.

### Calcium imaging

Calcium imaging was performed as previously described with some modifications (Okumura *et al*., 2022). In brief, cells were plated onto 96-well black-wall plates (#655090; Greiner Bio-One, Kremsmünster, Austria). After overnight incubation, the proteins were expressed by treatment with Dox. After 24 h, the cells were washed with PBS once, and incubated with a recording buffer (20 mM HEPES pH 7.4, 115 mM NaCl, 5.4 mM KCl, 1.8 mM CaCl_2_, 0.8 mM MgCl_2_, 13.8 mM _D_-Glucose, 1.25 mM probenecid) containing 2 μM Fluo 4-AM (#F311; Dojindo, Kumamoto, Japan) for 30 min. Probenecid, an organic anion transporter inhibitor, is often used for calcium assay to prevent the leakage of the loaded dye into the extracellular environment, which allows a stable amount of loaded dye in the cells during the experiments. The cells were then washed with PBS once and were incubated with the recording buffer for 30 min. For the calcium imaging, fluorescence images were captured every 2 s for 6 min using fluorescence microscopy (#AF6000-DMI6B; Leica Microsystems, Wetzlar, Germany) (objective lens; HCX PL FLUOTAR 20×/0.40 CORR PH1). The cells were stimulated with ATP solution by manual addition to the wells at the time point of 10 seconds. The data was measured from four independent wells. The captured fluorescence images were then analyzed by Fiji. 100 cells were randomly chosen from each image (total: 400 cells from four wells), and the center of each cell (8 × 8 pixels) was selected as a representative area of the cell. The Fluo-4 fluorescence intensity at 0 seconds was defined as F0, and the relative Fluo-4 intensity at each time point for 6 min was calculated as F/F0. The analyzed data (400 cells in each cell line) were used for the calculation of maximum AUC.

### Statistical analysis

The collected data were analyzed and graphically presented using GraphPad Prism7 (GraphPad, La Jolla, CA, USA). The statistical significances were represented as follows: \**P* < 0.05; \*\**P* < 0.01; \*\*\**P* < 0.001; \*\*\*\**P* < 0.0001. n.s.: not significant.

## Acknowledgments

We wish to thank Dr. Naonobu Fujita from Tokyo Institute of Technology for critical comment on our study. We also thank to Smart-Core-Facility Promotion Organization in TUAT for the technical support on the use of confocal microscopy and Institute of Global Innovation Research in TUAT.

## Data availability

The datasets generated during and/or analyzed during the current study are available from the corresponding author on reasonable request.

## Author contributions

Conceptualization: T.K., A.N.; Methodology: T.K.; Validation: T.K., C.J.; Formal analysis: T.K., C.J.; Investigation: T.K., C.J., W.O., N.O., T.Takao; Resources: T.K., C.J., W.O., H.S., H.M., T.Takagi, N.O., T.Takao; Data curation: T.K., C.J., W.O., N.O., T.Takao; Writing - original draft: T.K.; Writing - review & editing: N.O., T.Tonozuka, A.N.; Visualization: T.K., C.J., W.O. N.O.; Supervision: A.N.; Project administration: A.N.; Funding acquisition: T.K., A.N.

